# The trade-off between color and size in lizards’ conspicuous tails

**DOI:** 10.1101/2021.03.29.437563

**Authors:** Raiane dos Santos Guidi, Vinicius de Avelar São-Pedro, Holda Ramos da Silva, Gabriel Correa Costa, Daniel Marques Almeida Pessoa

**Affiliations:** Laboratory of Sensory Ecology, Department of Physiology & Behavior, Federal University of Rio Grande do Norte, Natal, RN, Brazil; Center of Natural Sciences, Federal University of São Carlos - *Campus* Lagoa do Sino, Buri, SP, Brazil; Department of Biology and Environmental Sciences, Auburn University at Montgomery, Montgomery, AL, USA

**Keywords:** Chromatic contrast, Color vision, Predator-prey interaction, Prey size, Tail autotomy

## Abstract

A tail of conspicuous coloration is hypothesized to be an advantageous trait for many species of lizards. Predator attacks would be directed to a non-vital, and autotomizable, body part, increasing the chance of survival. However, as body size increases it also increases the signaling area that could attract predators from greater distances, increasing the overall chance of predation. Here, we test the hypothesis that there is a trade-off between tail color and size, affecting predation probabilities. We used plasticine replicas of lizards to study the predation patterns of small and large lizards with red and blue tails. In a natural environment, we exposed six hundred replicas subjected to the attack of free-ranging predators. Large red-tailed models were attacked more quickly, and more intensely, by birds. Mammals and unidentified predators showed no preference for any size or colors. The attacks were not primarily directed to conspicuous tails when compared to the body or the head of our replicas. Our study suggests that red color signals in large lizards could enhance their detection by visually oriented predators (i.e., birds). The efficacy of conspicuous tails as a decoy may rely on associated behavioral displays, which are hard to test with static replicas.

**Highlights:**

1. The roles of blue and red tails as decoys were not corroborated.
2. Tail color and size interact while influencing predation rates.
3. Larger red-tailed lizards are more prone to be attacked by birds.
4. The benefit of having conspicuous tails appears to decrease as body size increases.

## 1. Introduction

Over evolutionary history, prey has developed several defense strategies to maximize survival rates in encounters with predators (Dawkins and Krebs, 1979). Predators play a strong evolutionary pressure over preys modifying their phenotypes (Castilla et al.,1999; Losos et al., 2004). In lizards, some of these strategies include escaping (Cooper, 2003; Hawlena, 2009; Schall and Pianka, 1980), deimatic behavior (Sherbrooke and Middendorf, 2001; Shine, 1990), morphological adaptations (Losos et al., 2002), tail autotomization (Bateman and Fleming, 2009), and cryptic or conspicuous color patterns (Fresnillo et al., 2015a; Stuart-Fox et al., 2004). Frequently, the same lizard species can combine two or more of these strategies in its predator avoidance repertoire (Pianka & Vitt, 2003; McElroy, 2019).

The use of color patterns as a defense strategy includes cryptic coloration, which conveys camouflage against dull backgrounds, decreasing the probability of detection by predators (Macedonia et al., 2004; Stuart-Fox et al., 2004). In contrast, conspicuous coloration can be used to frighten (Badiane et al., 2018) and discourage (Hasson, 1991) predators, or to redirect attacks to a non-vital region of the prey’s body, usually the tail (Bateman et al., 2014; Castilla et al., 1999; Fresnillo et al., 2015a; Murali and Kodandamaraiah, 2016; Ortega et al., 2014; Watson et al., 2012; Wilkinson, 2003). Such redirection can occur by distinct mechanisms, depending on whether the conspicuous coloration is related to a longitudinal striped body (Murali and Kodandamaraiah, 2016, 2017) or a colorful tail (Bateman et al., 2014; Fresnillo et al., 2015a).

Many lizard species from several unrelated families have tails with conspicuous coloration, which may be blue, green, or red, in contrast to the usually cryptic body-color (Murali et al., 2018). Although there are alternative hypotheses to explain the evolution of conspicuous tails (see Belliure et al., 2018; Clark and Hall, 1970;), several studies proposed that they can act as an effective decoy, directing predators’ attacks to the tail (Bateman et al., 2014; Castilla et al., 1999; Fresnillo et al., 2015a; Nasri et al., 2018; Ortega et al., 2014; Watson et al., 2012). Murali et al. also (2018) found that colorful tails in lizards are associated with diurnal species, suggesting that this trait has been selected against visually oriented predators. Nonetheless, it is still unknown to what extent predator-based selection has driven the evolution of color variation in lizards (McElroy, 2019).

Although conspicuous tails have received increasing attention in the last decades, there are still important questions to address on this topic. One of the most intriguing issues is the occurrence of this trait predominantly in small-bodied lizards (Pianka & Vitt, 2003). Even in some moderate-sized species, colorful tails occur mostly in juveniles, fading as the lizard approaches the minimal size for sexual maturity, usually around 40 mm of snout-to-vent length (Bateman et al., 2014; Castilla et al., 1999; Hawlena et al., 2006; Ortega et al., 2014). A possible explanation for this pattern links coloration to foraging mode, suggesting that colorful tails are only advantageous in juveniles due to their riskier behavior (Hawlena et al., 2006; Nasri et al., 2018). Such explanation does not consider that conspicuous tails persist in adult small-bodied lizard species (e.g., *Micrablepharus* spp. and *Vanzosaura* spp.), however, despite ontogenetic changes in their behavior.

In this study, we present the results of predation experiments using lizard replicas placed in the field. Our main goals were to test 1) the effectiveness of different tail colors as decoy and 2) the possible trade-off between tail color and body size in predation avoidance. We hypothesize that there may be a trade-off between tail size and its use as a colorful decoy for predator attack. While colorful tails would be beneficial for small lizards, the increase in body (tail) size would enhance the range of the color signal, potentially attracting more predators and becoming disadvantageous. We also hypothesize that tails reflecting longer wavelength colors (e.g., red) attract more predators than those exhibiting short wavelengths (e.g., blue), since shorter wavelengths suffer more Rayleigh scattering than longer wavelengths, being transmitted for shorter distances (Bradbury & Vehrencamp, 2011). On one hand, if the decoy hypothesis is correct, we expect that red-tailed and blue-tailed lizards will be attacked more frequently in the tail when compared to brown-tailed replicas. On the other hand, if the trade-off hypothesis is correct, large lizards with colorful tails (blue and red) should be more attacked than their smaller counterparts.

## 2. Material and methods

### 2.1 Lizard replicas

We hand-made a total of 600 lizard replicas from non-toxic white plasticine, in two different sizes, 300 large replicas (130 mm, total length) and 300 small replicas (60 mm, total length). Replicas were coated with non-toxic paint to resemble some of the color patterns exhibited by lizards of the study region (Figure 1S). Therefore, all replicas had brown dorsum and black flanks, but varied in tail color, which could be conspicuous (blue or red) or cryptic (brown), giving us six different experimental treatments that varied in color and size: large blue-tailed replicas, small blue-tailed replicas, large red-tailed replicas, small red-tailed replicas, large brown-tailed replicas, and small brown-tailed replicas (Figure 1).

**Figure 1.**
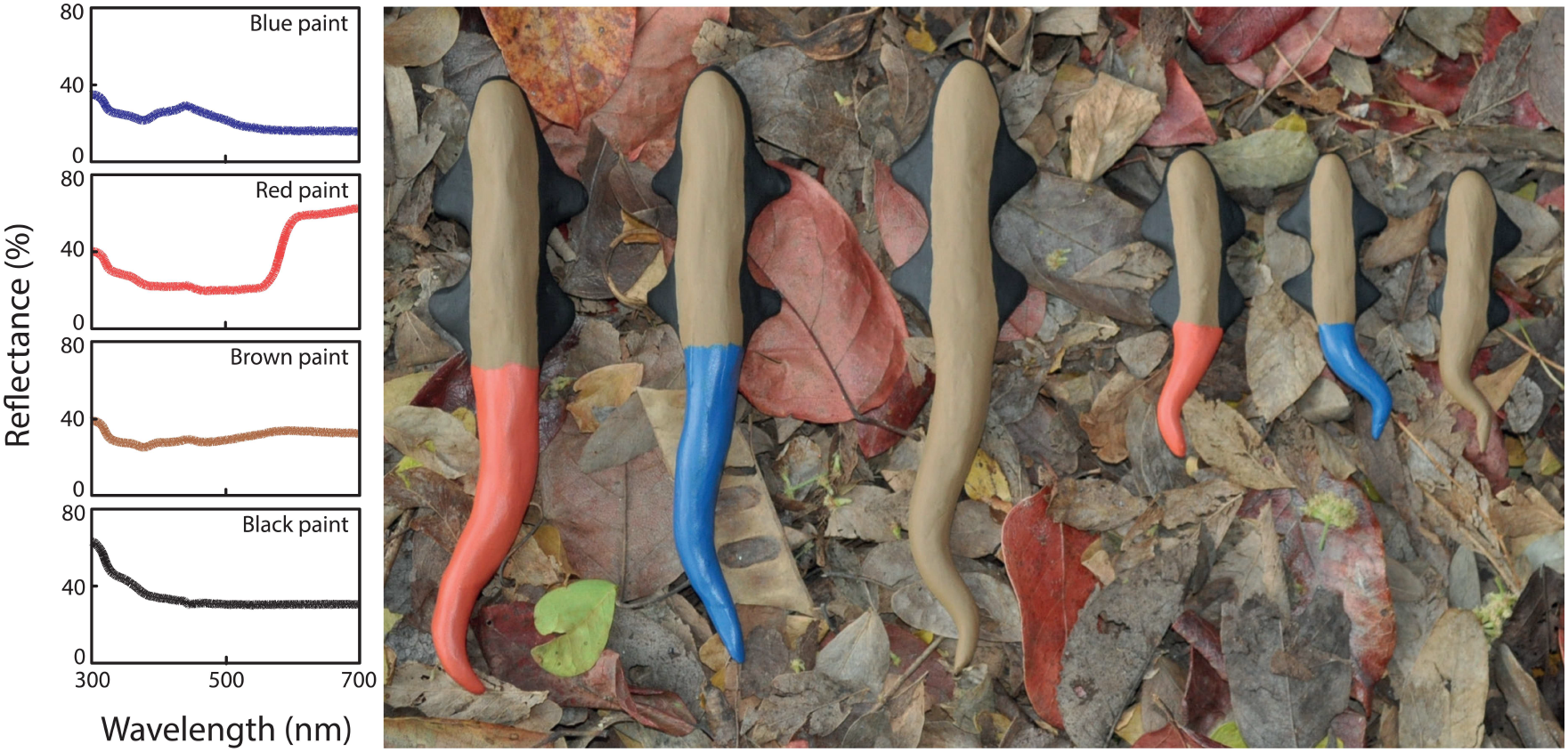
Six types of treatments used in our experiments. From left to right: large red-tailed replica, large blue-tailed replica, large brown-tailed replica, small red-tailed replica, small blue-tailed replica, and small brown-tailed replica. Small panels indicate the reflectance spectra of each kind of paint used to coat our replicas.

### 2.2 Predation experiments

Our research protocol was approved by the Ethics Committee on The Use of Animals of our institution (protocol 075/2015) and is following Brazilian law. It complies with ARRIVE guidelines and was carried out following the U.K. Animals (Scientific Procedures) Act, 1986, and associated guidelines, EU Directive 2010/63/EU for animal experiments.

Field experiments were carried out in *Parque Estadual da Dunas* (5° 48’ S; 35° 11’ W), an Atlantic Forest reserve located in Natal, the capital city of Rio Grande do Norte state, northeastern Brazil. Two small lizard species, with conspicuous tails, can be found in the state (Figure S1): the blue-tailed *Micrablepharus maximiliani* (Reinhardt and Lutken, 1861), which inhabits the very locality of the experiment, and the red-tailed *Vanzosaura multiscutata* (Amaral, 1933), that occurs in surrounding semiarid areas (Freire, 1996).

We conducted four repetitions of the same predation experiment, in 2017 (March, June, November) and 2018 (March). Each experiment consisted of placing 150 lizard replicas (25 of each experimental treatment) in the field for 20 days and recording predation marks left in the plasticine. Replicas were placed along a non-linear trail, at every five-meter interval, in such a way that their position could be alternated between the trail’s edge and forest interior (Figure S2). Replicas of different treatments were randomly assigned to each available position. Substrate did not vary along the trail and consisted of leaf litter over light brown sand. To prevent any predator from carrying the replicas away, each of them was attached to the nearest vegetation by a transparent fishing line, which was anchored to the bottom of the replica and remained hidden under the leaf litter.

Inspections for predation marks were made every two days. We recorded the following details if a replica had been attacked: to which treatment the attacked replica belonged; the type of mark left by the predator (bird peck, mammal bite, or unidentified predator); and in which body region the attack was launched (tail or body, which included head, trunk, and limbs). Marks left by ants (bite marks) were distinct and not counted as predator attacks (Shepard, 2007). Upon attack, replicas were removed from the field site and were not replaced, to avoid biased attacks made by accustomed predators (Marshal et al., 2015).

### 2.3 Spectrometry and visual modeling

To help us interpret the results of the experiment, we used visual modeling to understand how the main predators (i.e., birds) perceived the colors of replicas against the natural background. So, we measured the reflectance spectra (i.e., coloration) of each painted plasticine used to craft our replicas (Figure 1), as well as the reflectance spectra of the natural background (i.e., leaf litter), by using a USB4000-UV-VIS spectrometer (Ocean Optics Inc., Dunedin, Florida) connected to a laptop computer running software SpectraSuite (Ocean Optics Inc., Dunedin, Florida). Through a bifurcated QR450-7-XSR optic fiber (Ocean Optics Inc., Dunedin, Florida), the spectrometer was connected to a DH-2000-BAL light source (Ocean Optics Inc., Dunedin, Florida) and a probe. Measurements were taken at a 45° angle and a constant distance of five millimeters from the tip of the probe. For calibration of our spectrometric system, we used a spectralon reflectance standard WS-1-SL (Ocean Optics Inc., Dunedin, Florida) as our white standard, and turned the light source off, obstructing the probe orifice with a black cloth, for determining our black standard. Spectra averaged to scan and boxcar were set at 10 and 5, respectively.

The color difference between our replicas and the natural background was modeled using “pavo 2” (Maia et al., 2019), an R package. We ran receptor noise limited (RNL) models (Vorobyev et al., 1998), which gave us chromatic contrast values, in just noticeable difference (JND) units, between the reflectance spectra of our replicas and the leaf litter. Because the leaf litter at the experiment site consisted of leaves of varying colorations, we contrasted the coloration of our plasticine replicas (e.g., brown, black, blue, and red patches) with thirty-four different leaf litter measurements, and considered how the visual system of avian predators would discriminate replicas and background based on color alone.

There is a consolidated view that small deviations in receptor sensitivities do not affect model results significantly, it is, therefore, possible to use the spectral sensitivities of closely related species in models, if those of the species in focus are not known (Olsson et al. 2018). Since no information on the visual systems of native avian predators is available, we have employed blue tits’ (*Cyanistes caeruleus*) parameters in our visual model, which has been considered as a good proxy for passerine vision. The tetrachromatic vision of *Cyanistes caeruleus* counts with four classes of cones with different spectral sensitivities: ultraviolet sensitive cones (UV), blue-sensitive cones (S), green-sensitive cones (M) and red-sensitive cones (L). For calculation of absolute quantum catches for each of cone type we ran *vismodel*, from “pavo 2 package” (Maia et al., 2019), with the following arguments: *visual* = “bluetit”; *achromatic* = “none”; *illum* = “forestshade”, since our experiments took place in an Atlantic Forest area; *trans* = “bluetit”; *qcatch* = “Q_i_”; *bkg* = “ideal”; *vonkries* = “false”; *scale* = “1”; *relative* = “false”. Chromatic contrasts between replicas and leaf litter were calculated by using *coldist*, from “pavo 2 package” (Maia et al., 2019), with the following arguments: *noise* = “neural”, since we were only interested in modeling color signals for strictly diurnal predators (e.g., birds); *achromatic* = “false”; *n* = c(1,1.9,2.7,2.7), representing blue tits’ retinal relative proportion of photopigments (UV: 1.0; S: 1.9; M: 2.7; L: 2.7; Hart et al. 2000); *weber* = “0.1”; *weber.ref* = “longest”; *weber.achro* = “false”.

We classified the crypticity of our replicas’ patches according to the chromatic contrast they exhibited against the leaf litter. Following Siddiqi et al., (2004), we adopted three levels of detectability: cryptic (ΔS < 1 JND), poorly detectable (1 JND ≤ ΔS ≤ 3 JND) and detectable (ΔS > 3 JND). The higher the chromatic contrast, the higher the color difference between a replica and its surroundings, favoring their detectability.

### 2.4 Statistical analyses

First, we generated Kaplan-Meier survival curves for each treatment and compared them using Log-rank (Mantel-Cox), Breslow (Generalized Wilcoxon), and Tarone-Ware tests. To check the frequency distribution of attacks between all six treatments, we used Pearson’s chi-squared test. For survival analyses, replicas showing marks on the body and/or tail were considered as attacked (n = 223, from a total of 600 replicas).

We also build generalized linear mixed-effects models with R package lme4 vs. 1.1 (Bates et. al 2015) for testing whether attacks directed to the tail or body (response variable) were related to lizard size, tail color, or the interaction of size and color. Four different generalized linear mixed-effects models considered the attacks from 1) all predators combined, 2) mammals, 3) birds, and 4) mammals and birds. Replicas’ individual IDs were entered as a random factor.

We also compared the chromatic contrast between leaf litter and each replicas’ colored patches, by running a Friedmann test, with Wilcoxon *post-hoc*.

## 3. Results

### 3.1 Behavioral data

Among all the 600 replicas placed in the field, 223 (37%) suffered some kind of predator attack. All six treatments were attacked at least once, and most of the replicas attacked were large blue-tailed (n = 46, 21%), small red-tailed (n = 41, 18%) and large red-tailed (n = 39, 17%) (Figure 2). Unidentified predators were responsible for 78% (n = 174) of the attacks, followed by birds (n = 32, 14%) and mammals (n = 17, 8%).

**Figure 2.**
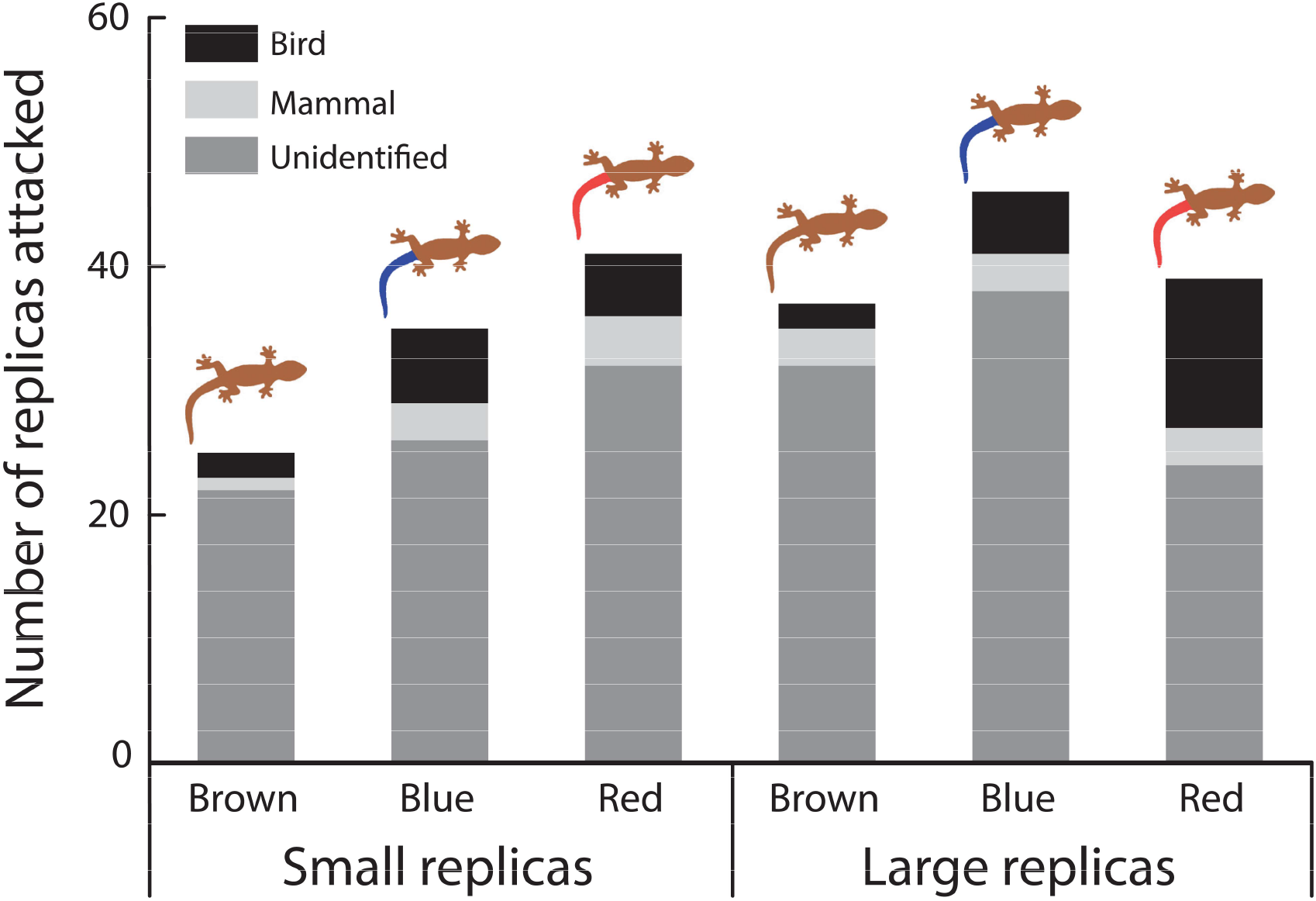
Number of attacks suffered by replicas of each treatment. Data from four repetitions are pooled. Total number of replicas used = 600.

Considering survival curves for the total number of replicas attacked (i.e., pooling data from all kinds of predators), we found no difference between treatments (Log-rank: df = 5, *P* = 0.08 / Breslow: df = 5, *P* = 0.12 / Tarone-ware: df = 5, *P* = 0.098). The same was true for the number of replicas attacked by mammals alone (Log-rank: df = 5, *P* = 0.807 / Breslow: df = 5, *P* = 0.807 / Tarone-ware: df = 5, *P* = 0.808) and by unidentified predators alone (Log-rank: df = 5, *P* = 0.141 / Breslow: df = 5, *P* = 0.272 / Tarone-ware: df = 5, *P* = 0.198). However, large red-tailed replicas were significantly more attacked by birds, when compared to replicas of other treatments (Log-rank: df = 5, *P* = 0.017 / Breslow: df = 5, *P* = 0.07 / Tarone-ware: df = 5, *P* = 0.01) (Figure 3).

**Figure 3.**
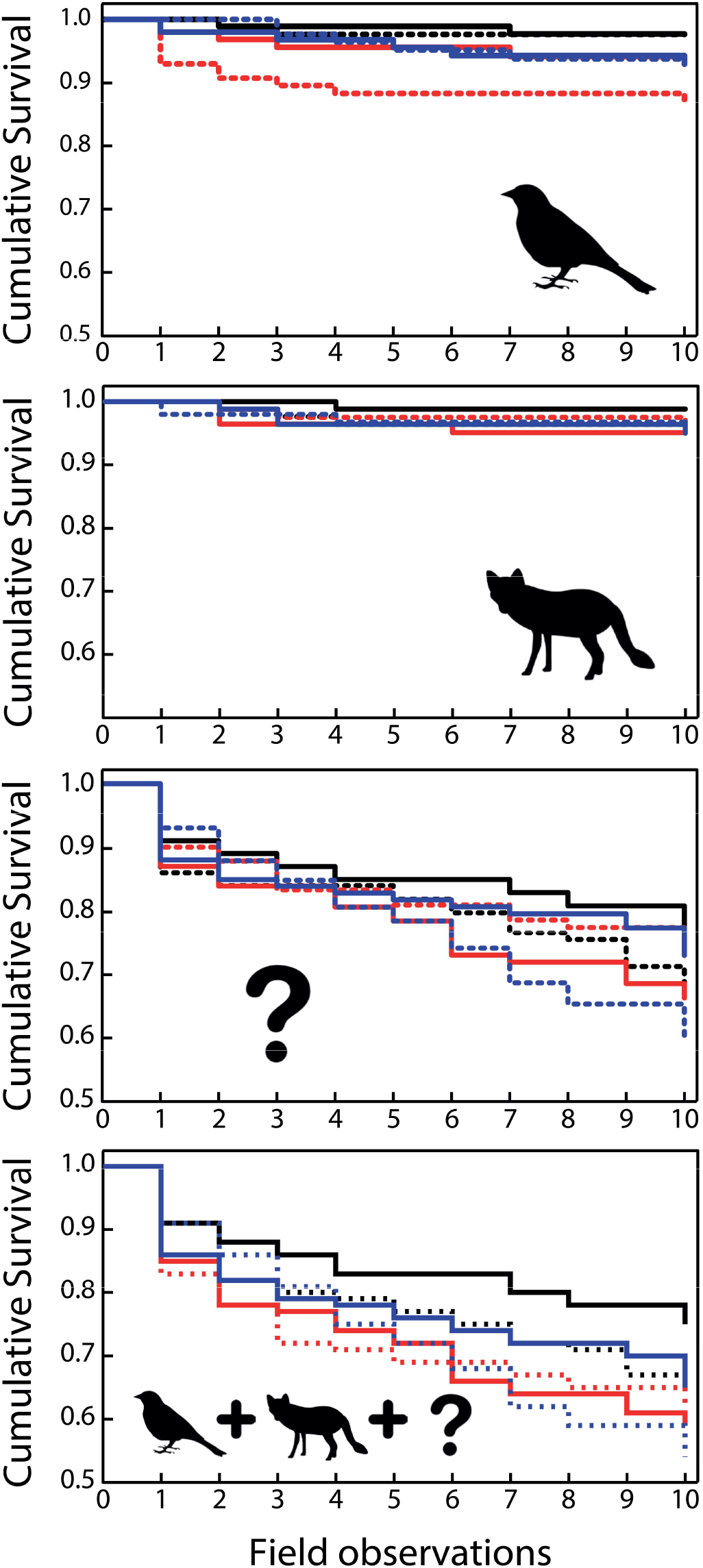
Kaplan-Meier survival plots for attacks performed by birds, mammals, unidentified predators (indicated by a “?” sign), and all kinds of predators pooled. Different treatments are indicated by different lines: small brown-tailed replicas = black solid line; large brown-tailed replicas = black dashed line; small red-tailed replicas = red solid line; large red-tailed replicas = red dashed line; small blue-tailed replicas = blue solid line; large blue-tailed replicas = blue dashed line. Data from four repetitions are pooled. Each repetition consisted of ten field observations, conducted throughout twenty experimental days. Total number of replicas used = 600.

The frequency of attacks between the six treatments was marginally significant (χ2 = 24.57, df = 15, *P* = 0.056). Values of adjusted residuals in the crosstab suggest that some frequencies are higher than expected by chance. More specifically, small brown-tailed replicas were less attacked than randomly expected (observed counts: 75; expected counts: 62.8; adjusted values: 2.8), large blue-tailed replicas suffered more unidentified attacks than expected (observed counts: 38; expected counts: 29; adjusted values: 2.2), while large red-tailed ones suffered more bird attacks than expected (observed counts: 12; expected counts: 5.3; adjusted values: 3.3) (Table S1).

Regarding our generalized linear mixed-effects models, that accessed the frequency of attacks directed to the tail or the body, we found no significant effect (Table S2) of color (p > 0.05), size (p > 0.05), or any interaction between these variables (p > 0.05). These results were consistent when we analyzed attacks performed by all kinds of predators, only by mammals, only by birds, and the attacks performed by mammals and birds (i.e., disregarding unidentified predators).

### 3.2 Visual modeling

Our visual model showed that, for visually-driven predators, such as passerines, every replicas’ body patches were expected to be detectable against the leaf litter (i.e., ΔS > 3 JND) (Figure 4). Blue and red tails were predicted to show equivalent levels of conspicuity (Z= −0.299, *P* = 0.765), although blue tails were predicted to be the most conspicuous of all body parts, contrasting significantly more from the leaf litter when compared to more cryptic body regions, such as black flanks (Z= −5.086, *P* < 0.0001) and brown dorsum/tails (Z=−5.086, *P* < 0.001). Red tails’ conspicuity was predicted to match that of black flanks (Z=−1.479, *P* = 0.139), and to differ from brown dorsum/tails’ chromatic contrast values (Z= −5.086 *P* < 0.0001). Brown dorsum/tails were also predicted to be more cryptic than black flanks (Z= −4.283, *P* < 0.0001).

**Figure 4.**
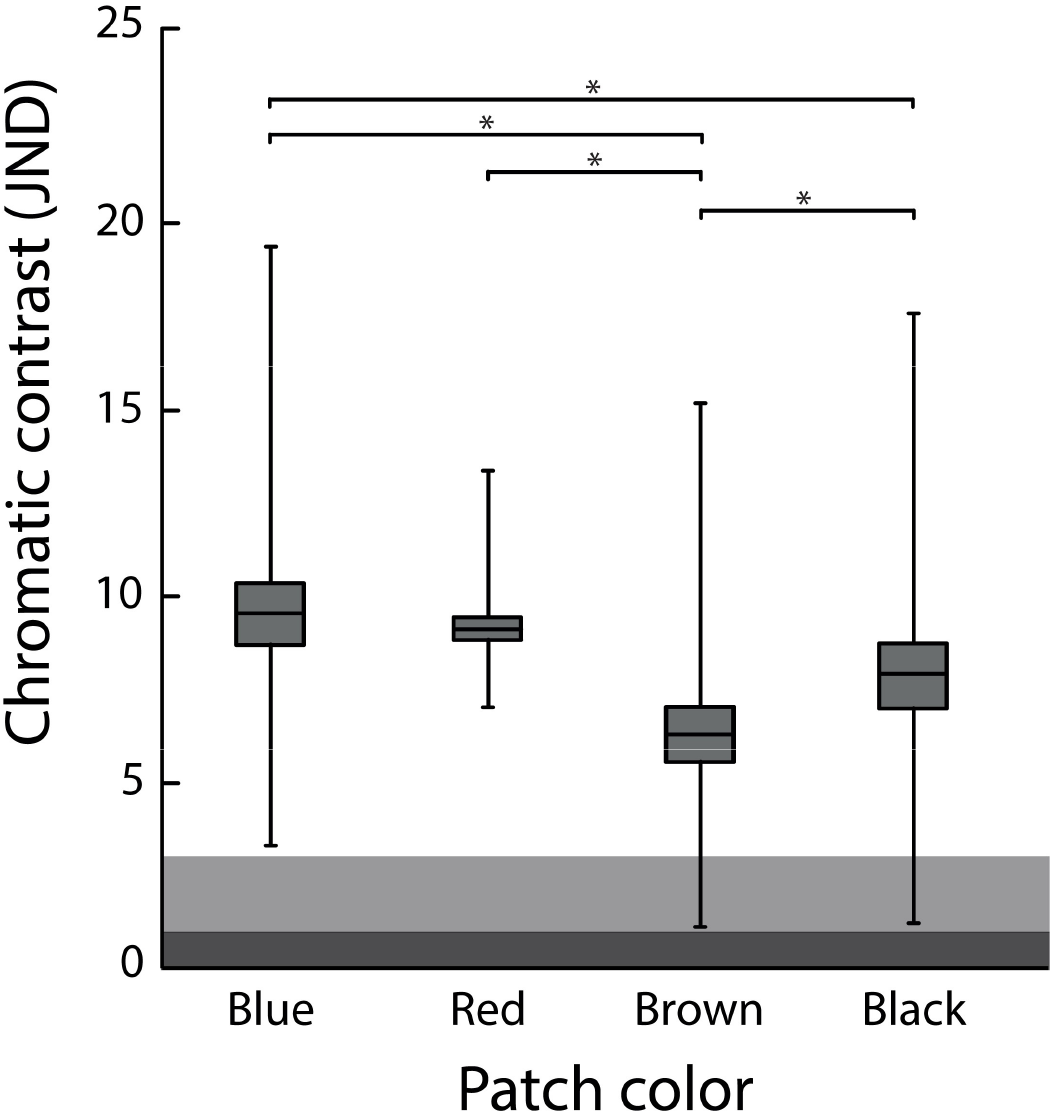
Chromatic contrast (ΔS) between replicas’ body patches (i.e., blue tail, red tail, brown tail/dorsum, and black flank) and leaf litter, according to the visual system of avian predators. Mean values are indicated by central horizontal black lines, SEM is limited by the boxes, while maximum and minimum values are indicated by whiskers. Detection thresholds are indicated by areas of different colors: the dark grey area refers to situations in which ΔS < 1 JND, the light grey area illustrates situations in which 1 JND ≤ ΔS ≤ 3 JND, while the white area indicates situations in which ΔS > 3 JND.

## 4. Discussion

Some of our results corroborate the hypothesis that there is a survival trade-off between body size and tail color in lizards. Although the results of our generalized linear mixed-effects were unable to indicate any effect of color or size on the frequency of attacks directed to replicas’ tails or body region, the hypothesis was supported by data from our survival analysis curves. Large red-tailed replicas were more frequently and rapidly attacked by birds than all other treatments, suggesting greater conspicuity of this kind of prey to visually oriented predators. Most birds are not just visually oriented but also rely primarily on color vision while foraging (Sherry, 2016). According to our visual model, birds should segregate blue and red tails from the background with the same aptitude. However, it is important to state that our visual model only took into consideration how a theoretical passerine visual system, under ideal conditions, would discriminate two objects based on color alone. Under field conditions many additional variables (e.g., outline, texture, brightness, glossiness, movement) gain importance in explaining the detection of colored targets, so modeling predictions only takes us to a certain point.

Although our visual model predicts blue and red tails to be equally conspicuous to birds, our behavioral data failed to confirm this prediction and showed, for example, large blue tails to be more cryptic than large red ones. This disparity between our theoretical predictions and our behavioral data might be, at least, partially explained through the effect of Rayleigh scattering, which disperses shorter electromagnetic waves through the air (e.g., blue color) more intensely than longer electromagnetic waves (e.g., red color), in such a way that blue signals tend to attenuate more vigorously and cannot be easily perceived at greater distances (Bradbury & Vehrencamp, 2011). Conversely, red signals travel longer distances and, consequently, might attract more predators, as shown by our predation experiment. When compared to smaller replicas, larger replicas produce stronger color signals, capable of enduring more attenuation, and propagating to longer distances. As birds take advantage of flight and perching on branches to scan the landscape for prey (Sherry, 2016), they are the predators that benefit most from long-distance signals. Birds are the most important predators of small to moderate-sized lizards and exert strong selective pressure on them (Pianka & Vitt, 2003), reinforcing the importance of such trade-offs in the evolutionary history of this group.

On one hand, our survival trade-off hypothesis seems to be adequate to explain the predominance of red tails in small lizards, since our results presume a size threshold from which having a red tail becomes more disadvantageous. On the other hand, our predation experiment showed that blue-tailed replicas were equally attacked by birds, regardless of body size, which does not explain the prevalence of blue-tailed lizards in small-bodied species. So, a still unanswered question is why blue tails do not persist in adult lizards of moderate sizes (e.g., several species of the family Scincidae). Pianka and Vitt (2003) hypothesized that these larger species lose their flashy tails to avoid the need for autotomy during adulthood when all energy must be saved to be used during the sparse reproduction episodes. In contrast, because tiny species, generally, produce more clutches per year, spending energy on regenerating a lost tail would not compromise much of their reproductive success (Pianka & Vitt, 2003). Yet, the same explanation might be applied to tails of other colors, including red tails. These hypotheses, as well as others mentioned before [e.g., increased movement hypothesis (Hawlena, 2009); aggression avoidance hypothesis (Fresnillo et al., 2015b)], are not mutually exclusive. None of these hypotheses alone seem to explain the occurrence of colorful tails in all their natural variation of size and colors. Only a more comprehensive work, which compiles the results of experimental studies and data on life-history traits of most lizard species with brightly colored tails, can elucidate the relative contribution of the mechanisms proposed in each hypothesis and the cases where they may have acted in synergy.

We did not find support for the hypothesis that conspicuous tails serve as decoys for predators. Our result contradicts similar studies with lizard replicas that endorse the efficacy of blue (Bateman et al., 2014; Watson et al., 2012), red (Fresnillo et al., 2015; Nasri et al., 2018), and green (Castilla et al., 1999) colors in redirecting predator attacks towards the tail. A possible explanation for that could be related to the type of predators recorded in each study. Most of these previous experiments based their conclusions on avian attacks solely and were conducted in temperate regions. In our study, conducted in a tropical forest site, even when we restricted our analyses to bird attacks, we found no preference for body or tail. We are aware that color itself may be not enough to redirect predators’ attacks to the tail. Indeed, several studies have suggested the importance of tail displays (e.g., lashing, wagging, waving) combined with conspicuous colors to effectively attract predator’s attention (Cooper & Vitt, 1985; Hawlena, 2009; Nasri et al., 2018). Despite the success of previous studies that used static replicas to corroborate the decoy hypothesis, we believe that movement stimuli can be decisive to direct the attack of some predators (Paluh et al., 2014). Perhaps our study did not corroborate this hypothesis because in our study area, unlike previous studies, there is a predominance of predators that depend on tail displays to unleash their attacks.

Regardless of inherent limitations, clay model experiments have proven to be an important tool for investigating predator-prey interactions (Bateman et al., 2017). While replicas seem to be convincing for birds (Paluh et al., 2015), there are still many doubts about their efficiency for other types of predators, such as mammals, snakes, and invertebrates, that might end-up prioritizing the use of non-visual sensory modalities when searching for prey. Indeed, in our experiment, unidentified predators accounted for most of the attacks (78%). These indistinct marks were likely left by arthropods, which are foragers mostly oriented by chemotactile cues (Greenfield, 2002). In this case, the damages in the replicas were randomly caused and their predominance in the body would be merely due to the greater volume of that part compared to the tail. To avoid misinterpretations, Bateman et al., (2017) recommend caution when attributing unidentified marks in clay replicas to predator attacks. Nevertheless, further investigations should be encouraged, to evaluate how replicas that mimic prey’s color and shape are effective in attracting predators and eliciting their response. Particularly, because it may be possible that non-avian predators have been underestimated by literature.

## 5. Conclusion

Here, we have experimentally shown, for the first time, the interaction of two important lizard morphological variables (color pattern and body size) as a predator avoidance strategy. Despite failing to confirm the effectiveness of conspicuous tails as decoys our experiment succeeded to show how a color signal that is beneficial for small-bodied and/or juvenile lizards may become disadvantageous for a larger-bodied and/or adult animal. Conspicuous tails have evolved independently, multiple times, among different groups of lizards (Murali et al., 2018). Understanding the evolutionary forces behind this amazing morphological trait, and its nuances, can be enlightening, not just for the study of lizards, but also for the understanding of predation ecology in general.

## 6. Acknowledgements

This study was financed in part by the Coordenacao de Aperfeicoamento de Pessoal de Nivel Superior – Brazil (CAPES) [Finance Codes 001 and 043/2012], which also provided a Post-Doc Scholarship to V.A.S.P.; and by Conselho Nacional de Desenvolvimento Cientifico e Tecnologico – Brazil (CNPQ) [Finance Codes 478222/2006-8, 25674/2009 and 474392/2013-9], that also granted two Scientific Initiation Scholarships for Undergraduate Students (R.S.G. and H.R.S.), as well as two Researcher Scholarships, to G.C.C. and D.M.A.P. We are also grateful to E. Oliveira, M.F. Erickson, S.C. Schirmer, G. Grimaldi, D.J.A. Silva, J.B.C. Neto, G. Mesquita, and R.T. M. Sales for helping with replicas’ crafting and field assistance; to IDEMA-RN (Nucleo de Gestao de Unidades de Conservacao) for allowing our research to be carried out in Parque Estadual das Dunas (Natal - RN); and to W. Silva for kindly providing the photos used in our Supplementary Material.

## Notes

Declarations of interest: none.

### Competing Interest Statement

The authors have declared no competing interest.

