## Supplementary Material for "The trade-off between color and size in lizards’ conspicuous tails"

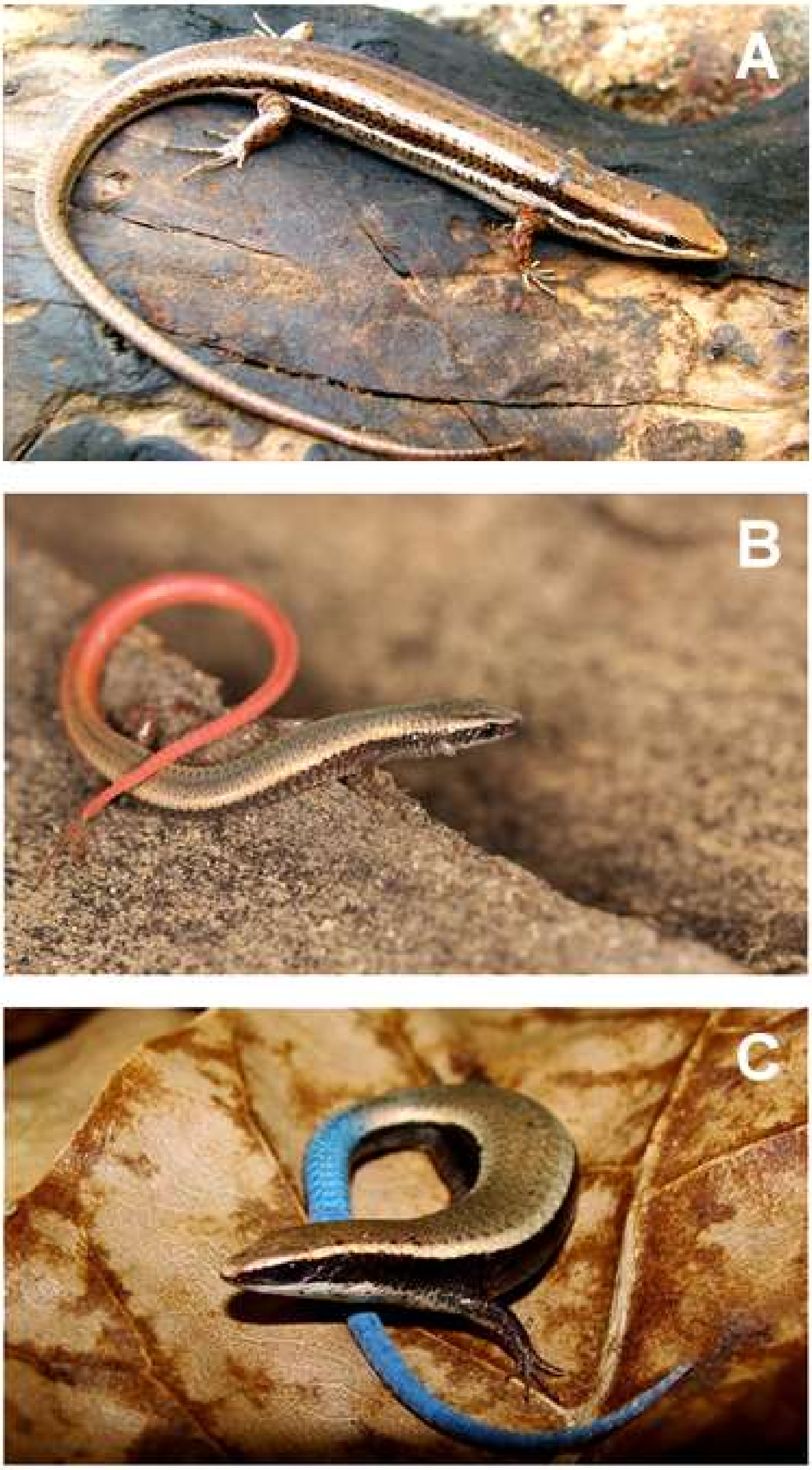


Figure S1. Lizard species that inspired us to create our replicas. A) *Brasiliscincus heathi*, showing a brown tail (Photo: Vinicius São-Pedro); B) *Vanzosaura multiscutata*, exhibiting a red tail (Photo: Willianilson Silva); and C) *Micrablepharus maximiliani* displaying a blue tail (Photos: Willianilson Silva).

### Supplementary material

Guidi, R.S., São-Pedro, V.A., Da Silva, H.R., Costa, G.C., Pessoa, D.M.A., 2021. The trade-off between color and size in lizards' conspicuous tails. Biorxiv - Animal Behavior and Cognition.

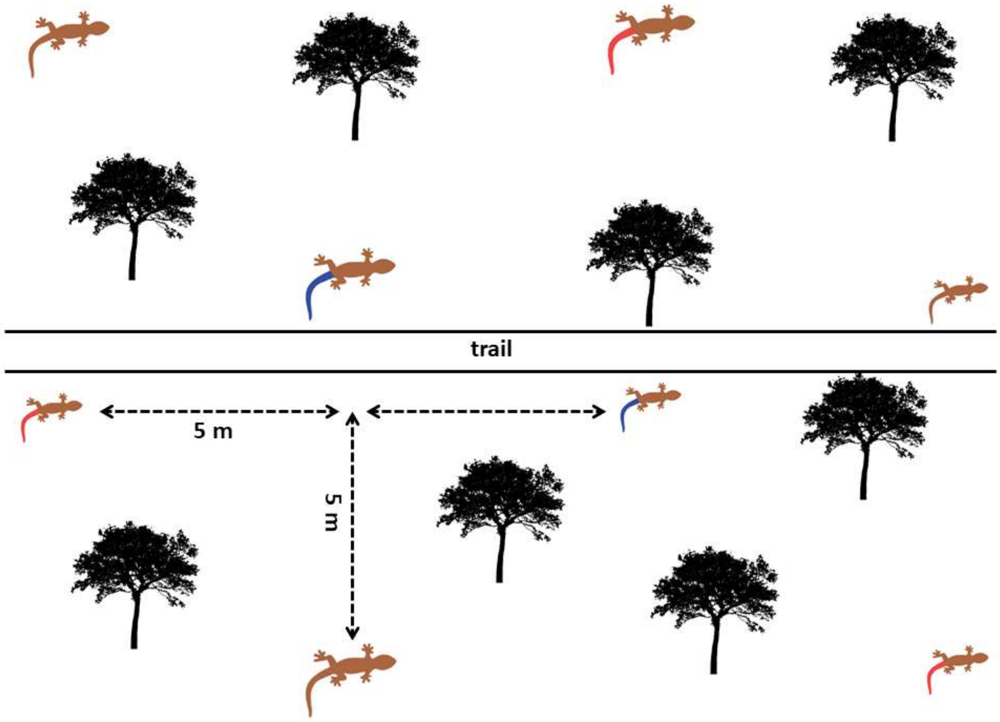


Figure S2. Schematic diagram showing how we placed our replicas along the experimental trail. Replicas of each treatment (color and size) were randomly allocated to each position.

### Supplementary material

Guidi, R.S., São-Pedro, V.A., Da Silva, H.R., Costa, G.C., Pessoa, D.M.A., 2021. The trade-off between color and size in lizards' conspicuous tails. Biorxiv - Animal Behavior and Cognition.

Table S1. Output from Pearson's chi-square analysis, with cross tabulation, concerning the distribution of predation events among treatments.

| Treatment | Not attacked | Mammal | Bird | Unidentified |
|---|---|---|---|---|
| **Small brown** | | | | |
| Observed counts | 75 | 1 | 2 | 22 |
| Expected counts | 62.8 | 2.8 | 5.3 | 29.0 |
| Adjusted values | **2.8*** | -1.2 | -1.6 | -1.7 |
| **Large brown** | | | | |
| Observed counts | 63 | 3 | 2 | 32 |
| Expected counts | 62.8 | 2.8 | 5.3 | 29.0 |
| Adjusted values | 0.0 | 0.1 | -1.6 | 0.7 |
| **Small red** | | | | |
| Observed counts | 59 | 4 | 5 | 32 |
| Expected counts | 62.8 | 2.8 | 5.3 | 29.0 |
| Adjusted values | -0.9 | 0.8 | -0.2 | 0.7 |
| **Large red** | | | | |
| Observed counts | 61 | 3 | 12 | 24 |
| Expected counts | 62.8 | 2.8 | 5.3 | 29.0 |
| Adjusted values | -0.4 | 0.1 | **3.3*** | -1.2 |
| **Small blue** | | | | |
| Observed counts | 65 | 3 | 6 | 26 |
| Expected counts | 62.8 | 2.8 | 5.3 | 29.0 |
| Adjusted values | 0.5 | 0.1 | 0.3 | -0.7 |
| **Large blue** | | | | |
| Observed counts | 54 | 3 | 5 | 38 |
| Expected counts | 62.8 | 2.8 | 5.3 | 29.0 |
| Adjusted values | -2.0 | 0.1 | -0.2 | **2.2*** |
| **Total** | | | | |
| Observed counts | 377 | 17 | 32 | 174 |

*Adjusted values that exceeded the limit of two (2.0) were considered as statistically significant.

### Supplementary material

Guidi, R.S., São-Pedro, V.A., Da Silva, H.R., Costa, G.C., Pessoa, D.M.A., 2021. The trade-off between color and size in lizards' conspicuous tails. Biorxiv - Animal Behavior and Cognition.

Table S2. Output from generalized linear mixed-effects models.

| | All Predators | | Mammals and Birds | | Mammals | | Birds | |
|---|---|---|---|---|---|---|---|---|
| Predictors | Odds Ratios | p | Odds Ratios | p | Odds Ratios | p | Odds Ratios | p |
| (Intercept) | 0.45 | 0.006 | 0.43 | 0.220 | 0.50 | 0.571 | 0.40 | 0.273 |
| Color [brown] | 1.62 | 0.239 | 4.67 | 0.164 | >10 | 1.000 | 1.25 | 0.880 |
| Color [red] | 1.36 | 0.471 | 1.48 | 0.639 | 1.00 | 1.000 | 1.67 | 0.605 |
| Size [small] | 1.12 | 0.800 | 2.33 | 0.346 | 8.00 | 0.210 | 1.00 | 1.000 |
| Color [brown] * Size [small] | 0.76 | 0.671 | 0.11 | 0.202 | 0.10 | 1.000 | 0.00 | 1.000 |
| Color [red] * Size [small] | 0.60 | 0.404 | 0.34 | 0.361 | 0.08 | 0.293 | 0.90 | 0.944 |
| Observations | 278 | | 61 | | 19 | | 42 | |
